# High dietary sucrose reveals distinct roles for the BBSome subunits BBS1 and BBS9 in *Drosophila* development

**DOI:** 10.64898/2026.11.23.768525

**Authors:** Sarolta Szentes, Vlad-Alexandru Toma, Alfréd Szabó, Emma Abigail Gardner, Kinga Kapusi, Katalin Farkas, Levente Kovacs

## Abstract

Insulin resistance and type 2 diabetes occur in Bardet–Biedl syndrome, and BBSome proteins regulate insulin receptor abundance at the cell surface in mammals. We asked how loss of two structurally related BBSome subunits, BBS1 and BBS9, affects development on a high-sucrose diet in Drosophila melanogaster. Sequence comparisons identified the seven-bladed β-propeller and γ-adaptin ear domains of human BBS1 and BBS9 in their respective fly orthologues. High sucrose (1 M) delayed pupariation in both single mutants, with the longest delay in flies lacking both subunits. The double mutant already showed a delay on standard sucrose (0.15 M). On high sucrose, its first pupae were recorded at 216 h after egg laying, whereas the last control and single-mutant pupae had formed by 168 h. Loss of BBS1 also reduced pupal volume and adult wing area on high sucrose. Both measurements remained close to control values in BBS9 mutants. These findings suggest overlapping contributions of BBS1 and BBS9 to developmental timing and a greater requirement for BBS1 to sustain growth under high dietary sucrose.

## Introduction

Bardet–Biedl syndrome (BBS) belongs to the disease family of inherited ciliopathies. The most common symptoms of BBS are retinal degeneration and renal disease, which are frequently accompanied by obesity and impaired glucose regulation (Melluso et al, 2023). In a UK cohort of 152 people with BBS, 24 had type 2 diabetes. Fasting glucose and insulin were also higher in the BBS group than in controls matched for age, sex and body mass index (Mujahid et al., 2018). The major group of proteins implicated in this syndrome assemble into the BBSome, an eight-subunit complex in mammals. The small GTPase ARL6 recruits the BBSome to membranes, where the complex recognizes cargo proteins and regulates their trafficking to cilia (Jin et al., 2010).

An emerging a link between BBS proteins and glucose regulation is demonstrated in studies of insulin receptor trafficking. Starks and colleagues found that depletion of BBS proteins reduced the abundance of insulin receptors at the plasma membrane. Fibroblasts from patients carrying the BBS1 M390R variant also had fewer receptors at the cell surface, and insulin elicited a weaker Akt response in tissues from BBS mutant mice (Starks et al, 2015). Deletion of Bbs1 in mouse hepatocytes likewise reduced surface insulin receptor levels and impaired hepatic insulin signaling (Rouabhi et al., 2023). These observations raise the question of whether the entire BBSome or just certain individual BBSome subunits elicit the response to excess dietary sugar. Further studies on animal models can contribute to address this question.

*Drosophila melanogaster* is one of the model organisms to be considered. In this laboratory model, insulin-like peptides act through a single insulin receptor (InR) to regulate cell growth (Brogiolo et al., 2001). High-sucrose feeding produces hyperglycemia in developing larvae, elevates circulating insulin-like peptide 2 and a reduces Akt response to insulin stimulation resembling the characteristic molecular mechanisms behind the type 2 diabetes (Musselman et al., 2011). A study by Pasco and Léopold also found that this insulin resistant larvae pupariated later and that the resulting adult males weighed less (Pasco and Léopold, 2012). Additionally, these studies link HSD-induced peripheral insulin resistance to growth inhibition, showing that larvae and resulting pupae are strikingly smaller than controls because *Drosophila* IIS cannot uncouple metabolic regulation from systemic growth. Thus, pupal volume is a readout of the animal’s sugar metabolism, and the volume attained at pupariation is established from pupal length and diameter (Delanoue et al., 2010). Adult wing area provides a complementary measure of organ growth. Timing and size can respond separately to changes in insulin signaling.

Moreover, reaction to InR activity during development is time sensitive. Reducing InR activity before larvae reach the critical size for metamorphosis prolongs development, while limiting it later limits final body and wing size (Shingleton et al., 2005).

Six out of eight BBSome subunits are conserved in Drosophila, including BBS1 and BBS9, being the only two subunits having 7-bladed beta propeller and gamma-adaptin ear domains (Klink et al, 2020). Two other BBSome subunits, BBS7 and BBS7 are reported to contain these domains in humans, however their orthologues are not found in Drosophila. In our earlier work, we demonstrated that both BBS1 and BBS9 single mutants were viable and fertile under standard culture conditions, but males lacking both subunits had reduced fertility (Szentes et al., 2026).

This genetic interaction indicates some degree of functional complementation and led us to ask whether BBS1 and BBS9 also make overlapping contributions to development under conditions to induce insulin resistance. Here, we compare the conservancy of their domain organization and examine how changing dietary sucrose affects pupariation and growth. The two mutations together produced the longest delay in pupariation. Among the single mutants, loss of BBS1 produced the clearest reduction in pupal volume and adult wing area corresponding to the readout of a diabetic condition.

## Materials and Methods

### Drosophila stocks

Drosophila stocks were maintained on standard yeast-cornmeal-sucrose medium at 25 °C unless otherwise indicated. Details of generation of BBS1 and BBS9 deletion null alleles were described in Szentes et al, 2026. In brief, both alleles were generated by targeting the 5’ and 3’ end of coding sequences simultaneously by CRISPR/Cas9 resulting in removal of the entire coding region. Standard *w*^*1118*^ line was used as a control in each experiment.

### Bioinformatical analyses

Amino acid seqeuences of human BBS1 and BBS9 were downloaded from UniProt database. For Drosophila counterparts we used the protein coding sequences reported in FlyBase.

Global and local, domain specific comparison were performed by using EMBL-EBI GGSEARCH2SEQ Pairwise Sequence Alignment tool.

### Sucrose feeding and measuring developmental time

Standard yeas-cornmeal-sucrose medium were prepared, however the concentration of added sucrose was modified according to the low (0.02 M), normal (0.15 M) and high (1 M) sucrose content. Control, *BBS1*^*null*^ homozygote, *BBS9*^*null*^ homozygote and, *BBS9*^*null*^; *BBS1*^*null*^*/CyO::TM6B* double mutant heterozygote adult pairs (approx. 60 pairs each) were kept in egg laying chamber for synchronous embryo collection on agar plates. Forty freshly hatched, 0-2 hours synchronized L1 larvae were collected into each vial with low, normal or high sucrose medium. At least 5 parallel vials were established for each genotype and sucrose concentration in a single experiment. Each experiment were three times repeated using biological replicates. Vials were monitored in every 12 hours and number of pupae were recorded. Data were analyzed using Microsoft Excel and its embedded statistical analysis tools.

### Measuring pupal volume and wing surface area

For volumetric analysis, pupae were collected from vials into glass microscopic slides using a wet paintbrush. After pupae dried and stuck to the glass surface, they were images by an Olympus SZ51 stereomicroscope along with corresponding measuring references. Images were processed by ImageJ, length and diameter of each pupae were measured. Pupal volumes were calculated using the formula 4/3π(L/2)(l/2)^2^ (L, length; l, diameter) published in Delanoue et al. (2010).

For wing surface are measurement, wings of adults flies were dissected in 70% EtOH and transferred into 12 µl 50% glycerol on a microscope slide. Images were taken using an Olympus CX23 microscope and were processed in ImageJ. Wing area were calculated from the measured wing circumference.

## Results

### BBS1 and BBS9 characteristic domain arrangements are conserved in Drosophila

To identify structural elements that could underlie a shared function, we aligned Drosophila BBS1 and BBS9 with their human orthologue and mapped the domains using the mammalian BBSome structure described by Chou and colleagues (Chou et al, 2019). Fly and human BBS1 share 29.5% amino acid identity and 50.9% similarity over the full-length alignment. The corresponding values for BBS9 are 27.1% and 54.7% (Figure 1). In each protein, a seven-bladed β-propeller and a γ-adaptin ear (GAE) domain was present. Human and fly sequences sequence similarity within the GAE domain is 65% for BBS1 and 52% for BBS9 respectively. The GAE lies near the C terminus of BBS1. In BBS9 it is followed by an α/β platform and a helical region in both species. Senqeunces encoding the blades of the seven-bladed β-propeller domain in both species and in both proteins resembled a 60-80% conservation. This data indicates that in the potentially functional domains there is a higher degree of conservation that at the other part of the protein sequences. Because Drosophila lacks BBS2 and BBS7, BBS1 and BBS9 are the only fly BBSome subunits with this combination of propeller and ear domains (Klink et al., 2020).

**Fig 1.**
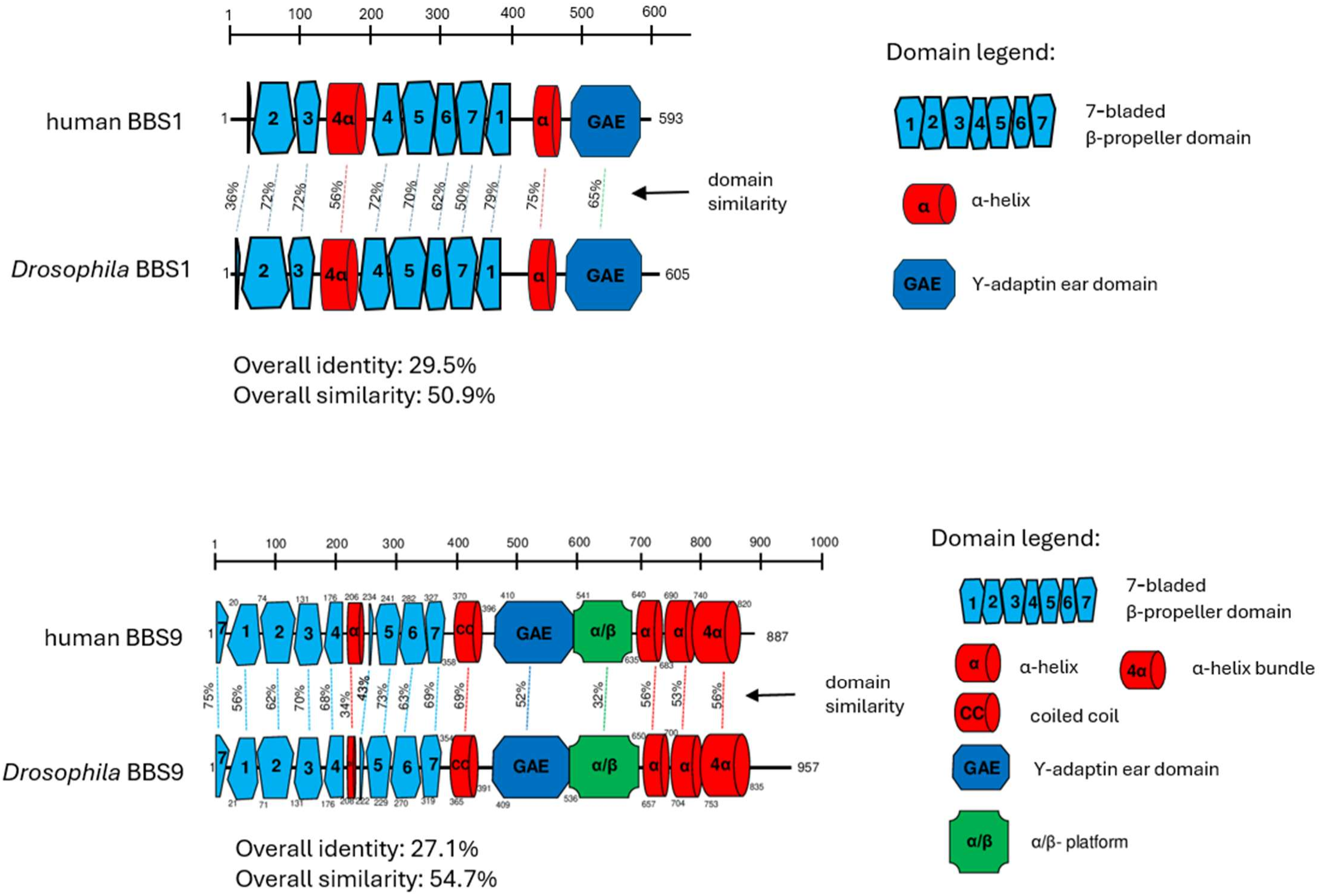
Sequence and structural homology between human and Drosophila BBS1 and BBS9 proteins. Human and fly BBS1 (upper panel) and BBS9 (lower panel) were compared by pairwise alignment using EMBL-EBI GGSEARCH2SEQ. The scales indicate amino acid positions. Dashed lines connect corresponding structural elements, with the percentage sequence similarity indicated alongside each line. Full-length sequence identity and similarity are given below each pair. The domain schematics follow Chou et al. 2019.

Notably, the gene encoding the BBS1 BBSome subunit is the most frequently mutated in Bardet-Biedl Syndrome patients, accounting over 20% of the cases (Maskova et al, 2026), however, the mutations in the structurally similar BBS9 are less frequent, in about 5% of all BBS cases (Xin-Yi et al, 2023). Their shared architecture and yet the differential contribution to the disease prompted us to ask how far one subunit might support development in the absence of the other in Drosophila.

### Combined loss of BBS1 and BBS9 delays pupariation even on standard sucrose

As high sucrose content of Drosophila food triggers insulin resistance, which is also a frequent symptom in BBS, we tested the sucrose sensitivity of the *BBS1*^*null*^ and *BBS9*^*null*^ alleles. The coding-region deletions were generated by CRISPR/Cas9 and their breakpoints verified by sequencing in our earlier study (Szentes et al., 2026). First-instar larvae were reared on low (0.02 M), standard (0.15 M) or high (1 M) sucrose medium. At 120 h after egg laying, pupae were visible on the wall of the high-sucrose control vial. In the representative vials shown for each single mutant, the walls were still free of visible pupae on the same diet (Figure 2A).

**Fig 2.**
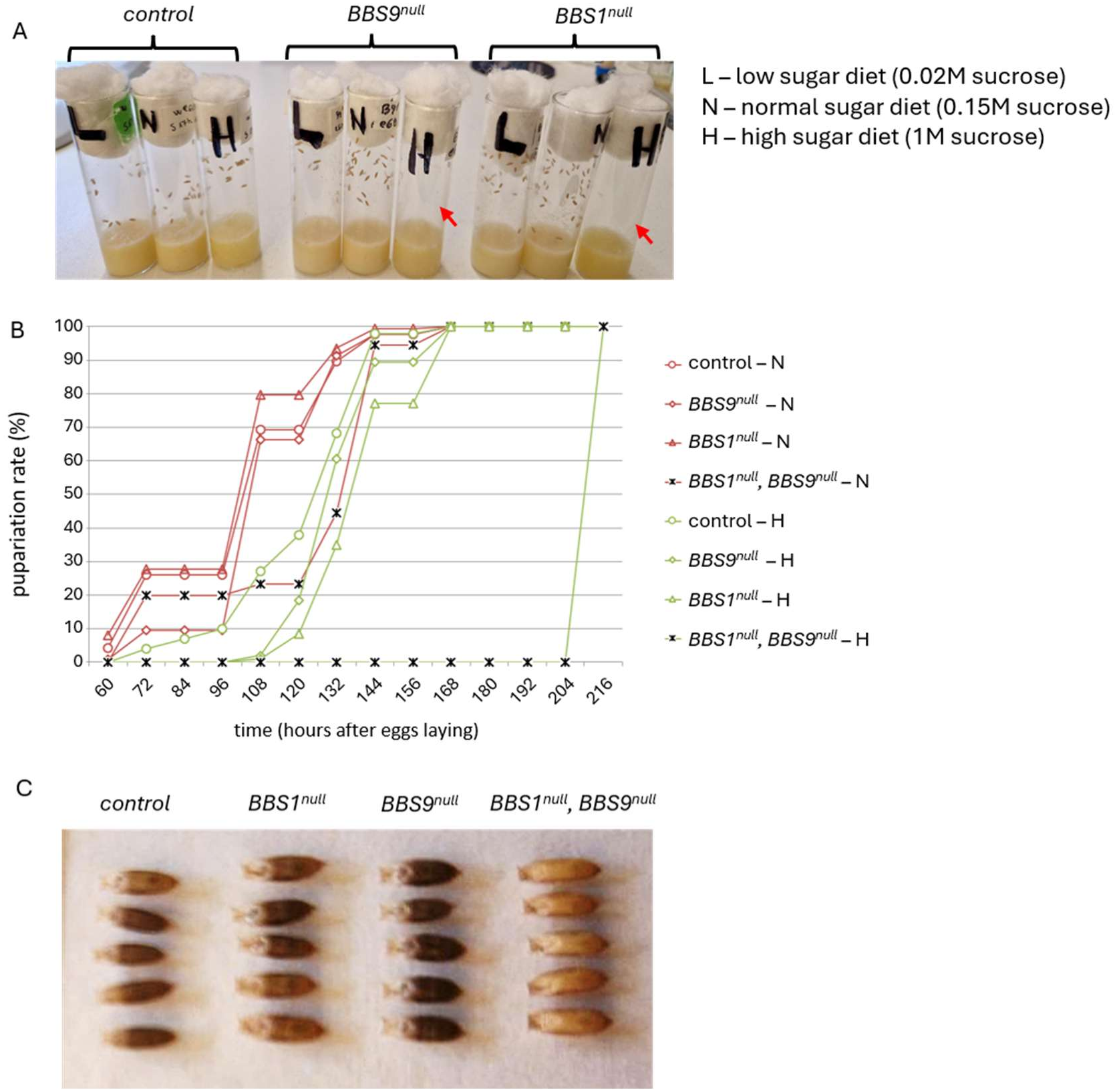
Sucrose sensitivity of BBS1 and BBS9 mutants. (A) Representative vials photographed 120 h after egg laying. Each vial received 40 first-instar larvae aged 0–2 h. L, N and H indicate low (0.02 M), standard (0.15 M) and high (1 M) sucrose, respectively. Red arrows mark high-sucrose single-mutant vials with no pupae visible on the wall. (B) Pupariation on standard and high sucrose. Pupae were counted every 12 h. For each genotype and diet, the percentage at each time point is expressed relative to the total number that ultimately pupariated. Five parallel vials were established per genotype. Plotted values are averages from three independent biological replicates. (C) Pupae collected 168 h after egg laying from synchronized cohorts reared on standard sucrose. The control and single-mutant pupae show the darker pigmentation of later pupal development. The double-mutant pupae are light brown.

We scored pupariation at 12-h intervals on standard and high sucrose. In this experiment we also included the testing the *BBS1*^*null*^-*BBS9*^*null*^ double mutant. On standard sucrose, approximately two thirds of the controls and BBS9 mutants and four fifths of the BBS1 mutants had pupariated by 108 h. Notably, the double mutant had a significant delay in development and only about one fifth of them pupariated by 108h timepoint.

As expected, high sucrose delayed pupariation in all genotypes. At 132 h, approximately 68% of the controls, 60% of the BBS9 mutants and 35% of the BBS1 mutants had pupariated on high sucrose (Figure 2B). By 168 h all three groups had reached their final pupal counts. However, no double-mutant pupae were recorded on high sucrose through 204 h. The first were scored at 216 h, two days after the control and single-mutant groups had completed pupariation. The baseline delay was also evident from the pigmentation of pupae collected the same time after egg laying. At 168 h on standard sucrose, control and single-mutant pupae had darkened late pharate adult stage, while double-mutant pupae were still early brown pupal stage (Figure 2C).

### High sucrose reduces pupal volume and adult wing area in BBS1 mutants only

To determine whether the delay in either single mutant was accompanied by reduced growth, we compared pupal volume across all three diets (Figure 3A). Mean pupal volume in BBS1 mutants was approximately 1.0 mm^3^ on both low and standard sucrose and showed a markedly decrease to about 0.7 mm^3^ on high sucrose. Contrarily, on the high-sucrose diet, pupal volume averaged approximately 1.1 mm^3^ in controls and 1.0 mm^3^ in BBS9 mutants.

**Fig 3.**
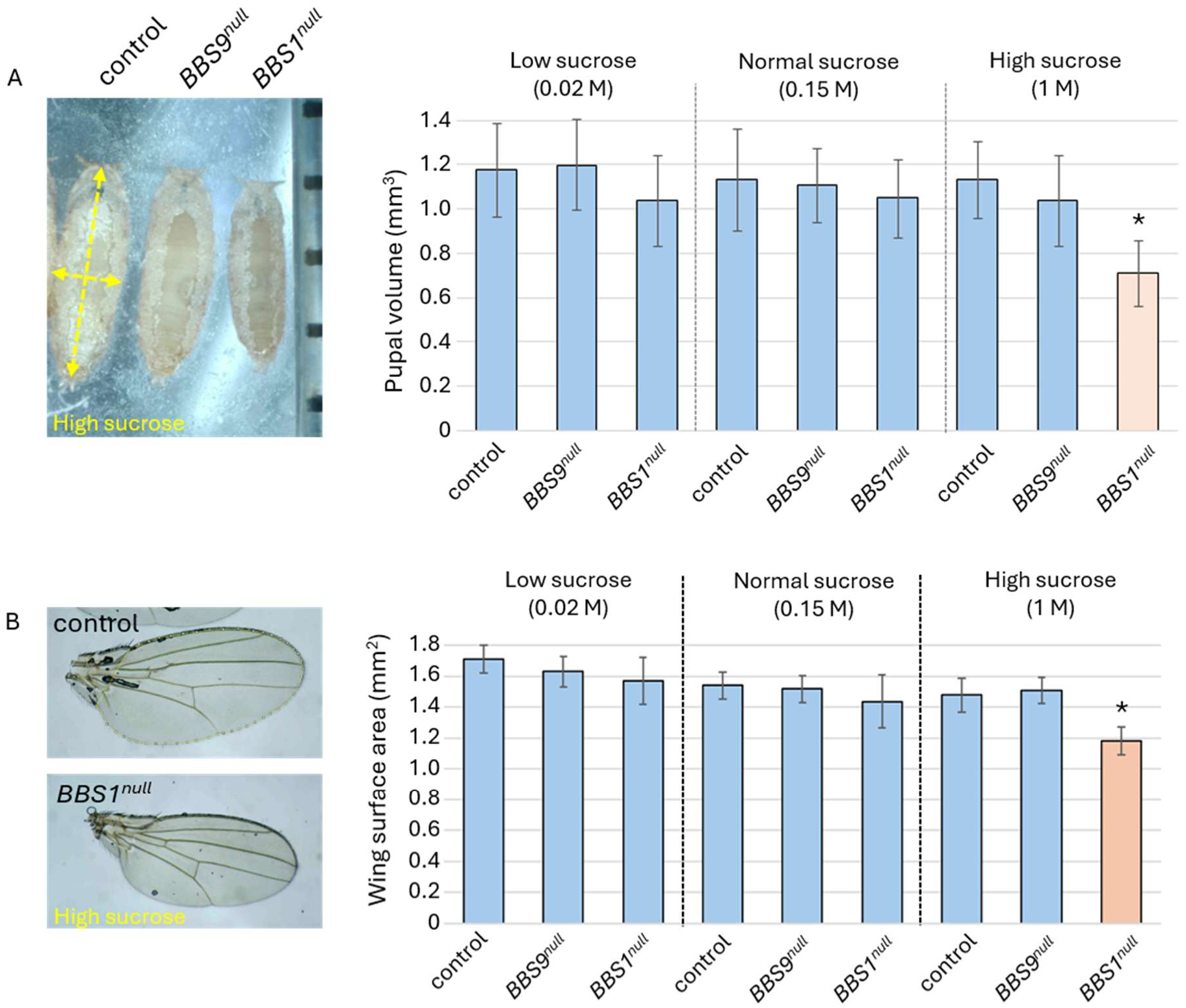
High sucrose concentration leads to small size in *BBS1*^*null*^ mutant. (A) Representative ventral views of pupae reared on high sucrose and mean pupal volume on each diet. Dashed arrows indicate the length and diameter used to estimate volume as described by Delanoue et al. (2010). At least 100 pupae were imaged per genotype and condition. (B) Representative adult wings from high-sucrose control and *BBS1*^*null*^ flies and mean wing area on each diet. Wings were mounted in 50% glycerol, their outlines traced, and their areas measured in ImageJ. At least 100 wings were imaged per genotype and condition. Bars in both panels show means and error bars indicate standard deviations. * p < 0.05.

We next asked whether the difference in size was also evident also in adult, quantitatively indicated by wing surface (Figure 3B). Mean wing area in BBS1 mutants fell from approximately 1.4 mm^2^ on standard sucrose to 1.2 mm^2^ on high sucrose. On high sucrose, wings of both controls and BBS9 mutants averaged about 1.5 mm^2^. Thus, the growth deficit in high-sucrose BBS1 mutants was detectable both at pupariation and in an adult organ, whilst the absence of BBS9 subunit did not manifest in such phenotype.

## Discussion

Reduced cell-surface expression of the insulin receptor offers one explanation for insulin resistance in Bardet–Biedl syndrome (Starks et al., 2015). We used the growth response to dietary sucrose in Drosophila to ask how two BBSome subunits contribute to development under these conditions. The strongest effect on developmental timing occurred when both BBS1 and BBS9 were absent. These animals were delayed even on standard food, and on high sucrose the other genotypes had finished pupariating before the first double-mutant pupae appeared. The two single mutants differed more clearly in growth. In BBS9 mutants the modest delay on high sucrose was accompanied by pupal volumes and adult wing areas close to those of controls. In BBS1 mutants, pupae and wings were both smaller despite the additional time spent in larval development.

One interpretation is that the subunits can partially compensate for one another. BBS1 may sustain sufficient growth when BBS9 is absent, while BBS9 may be less able to sustain growth in the absence of BBS1 on high sucrose. The greater delay in the double mutant is consistent with both subunits contributing to developmental timing. Their domain organization offers a basis for considering functional overlap, although the human BBSome structure also shows distinct interactions. The β-propeller of BBS1 contacts BBS4 and provides a binding site for ARL6, while the corresponding propeller of BBS9 contacts BBS8 (Klink et al., 2020). These distinct interactions could give BBS1 a specific role in growth. Measuring size in the double mutant and testing how far each subunit, expressed separately at comparable levels, restores growth and pupariation would help distinguish this possibility from asymmetric compensation. The BBSome’s role in insulin receptor trafficking suggests a mechanism for the growth defect. In mice, deleting Bbs1 in hepatocytes impairs glucose handling and insulin signaling, while skeletal muscle deletion leaves glucose handling largely intact (Rouabhi et al., 2023). In the fly, a reduced pool of surface InR in the larval fat body or wing imaginal disc could likewise alter the growth response to dietary sugar. The stage at which such a defect occurs could help explain why the two mutants differ. InR activity influences developmental timing early in larval life and final body and wing size later in development (Shingleton et al., 2005). A requirement for BBS1 during the later growth period could account for the reduction in size specific to this mutant.

The relevance of these phenotypes to type 2 diabetes rests on whether the mutant tissues become insulin resistant. In high-sugar fly models, developmental delay and reduced size accompany elevated circulating sugars and diminished insulin responsiveness (Musselman et al., 2011).

Differences in food intake could also affect growth, as could changes in ecdysone signaling in the larval fat body (Delanoue et al., 2010). Measuring hemolymph glucose and insulin-stimulated Akt phosphorylation in stage-matched larvae would establish whether impaired glucose regulation accompanies the BBS1 growth defect. The single-mutant comparison offers a specific test of the trafficking model. If reduced receptor availability accounts for the difference in growth, high-sucrose BBS1 mutants should show a greater reduction in the fraction of InR at the cell surface than BBS9 mutants at the same developmental stage.

## Aknowledgement

We are grateful to Prof. David M. Glover for many advice and support. We are thankful to Orsolya Dózsa for technical help provided. This work was supported by PN-IV-P2-2_1-TE-2023-0123 grant from UEFISCDI office of Romanian Ministry of Education and Research to Levente Kovacs.

## Notes

### Competing Interest Statement

The authors have declared no competing interest.

